# *De novo* prediction of RNA-protein interactions with Graph Neural Networks

**DOI:** 10.1101/2021.09.28.462100

**Authors:** Viplove Arora, Guido Sanguinetti

## Abstract

RNA-binding proteins (RBPs) are key co- and post-transcriptional regulators of gene expression, playing a crucial role in many biological processes. Experimental methods like CLIP-seq have enabled the identification of transcriptome-wide RNA-protein interactions for select proteins, however the time- and resource-intensive nature of these technologies call for the development of computational methods to complement their predictions. Here we leverage recent, large-scale CLIP-seq experiments to construct a *de novo* predictor of RNA-protein interactions based on graph neural networks (GNN). We show that the GNN method allows not only to predict missing links in an RNA-protein network, but to predict the entire complement of targets of previously unassayed proteins, and even to reconstruct the entire network of RNA-protein interactions in different conditions based on minimal information. Our results demonstrate the potential of modern machine learning methods to extract useful information on post-transcriptional regulation from large data sets.

## 1 Introduction

RNA-protein interactions are fundamental in the regulation of gene expression. RNA-binding proteins (RBPs) are key in RNA splicing, processing, export, localization, and regulation of translation. Despite their importance, RNA-protein interactions are still relatively understudied, when compared with the DNA-protein interactions which are involved in the initiation and regulation of transcription. Many proteins with previously unsuspected RNA-binding properties are still being discovered, and more than 2000 human proteins have been experimentally determined to bind RNA (Brannan *et al*., 2016; Hentze *et al*., 2018; Liu *et al*., 2019). However, due to the difficulty of experimentally determining interactions between individual proteins and individual transcripts, the number of RBPs for which the identity of their interaction partners is known is much lower.

A major breakthrough in the study of RNA-protein interactions was the development of high-throughput techniques such as CLIP-seq (cross-linking and immunoprecipitation followed by next generation sequencing) (Licatalosi *et al*., 2008). CLIP-seq enables the isolation of RBPs and the fragments of RNA which are bound to them, much in the way that ChIP-seq is used to determine regions of DNA bound by transcription factor proteins. Thanks to improvements in the technology such as the development of the eCLIP protocol (Van Nostrand *et al*., 2016), huge numbers of RBP binding sites are being verified. Despite these advances, practical and conceptual hurdles mean that we are still very far from a comprehensive mapping of the network of RNA-protein interactions. First of all, such networks are intrinsically condition dependent (for example, simply because specific transcripts might be present or absent in different conditions). Secondly, the experimentally determined interactions are inevitably noisy, meaning that both false positives and false negative results are likely. Thus, there is a need for computational methods to complement experimental techniques by filtering noise and predicting interactions for new conditions as well as new RBPs. Here, we consider the problem of predicting RNA-protein interaction (RPI) pairs adopting a machine learning perspective, where a model is trained on currently validated interactions, using RNA and protein sequences as inputs.

Most current predictive methods focus on the narrower problem of predicting binding sites for a specific protein within RNAs, often combining sequence and secondary structure of the target transcript (Maticzka *et al*., 2014; Kazan *et al*., 2010; Alipanahi *et al*., 2015; Uhl *et al*., 2021). Due to the lack of large scale CLIP-seq datasets in the past, methods for predicting RNA-protein interaction pairs have only been trained on small datasets (Pan *et al*., 2019, Section 4.1). RPIseq (Muppirala *et al*., 2011) uses the sequence information of RNAs and proteins to predict interactions using SVM and random forests as classifiers. catRAPID omics (Agostini *et al*., 2013) uses the physiochemical properties of sequences to predict RNA-protein interactions on a genome-wide scale. Deep learning-based methods were also proposed (e.g. IPMiner (Pan *et al*., 2016), RPI-SAN (Yi *et al*., 2018), RPIFSE (Wang *et al*., 2019), RNAcommender (Corrado *et al*., 2016), and ELM* (Wang *et al*., 2018)) but due to data paucity they were not trained in an end-to-end way and usually relied on advanced feature engineering. As far as we can tell, all models for RNA-protein interaction prediction, such as RPIseq (Muppirala *et al*., 2011), IPminer (Pan *et al*., 2016), and recent models like NPI-GNN (Shen *et al*., 2021) have been designed for small datasets with a handful of proteins and targets (see Table S1). The only method we found that was developed with large datasets in mind was RNAcommender (Corrado *et al*., 2016) (which in the original paper was trained and tested on heterogeneous data from different experiments).

In this paper, we propose to exploit new, large-scale eCLIP datasets (Van Nostrand *et al*., 2020) to shift the problem of RNA-protein interaction prediction to the network level, i.e. attempting to predict the *whole network* of RPI in an organism in a particular condition in an end-to-end way. We use graph neural networks for predicting RPI, moving away from the paradigm of predicting the targets of a single protein towards leveraging whole network information. To achieve this, we curate a dataset of RNA-protein interactions in homogeneous conditions using the high-throughput CLIP-seq data generated as part of the ENCODE project (Van Nostrand *et al*., 2020) to train our models. We also show that the model can be used to predict the interactions for previously unseen proteins as well as transfer the knowledge to a network observed under different biological conditions. We achieve this by using the similarity between the sequence-based features of proteins to elicit the embedding for a previously unseen protein. The results show the superiority of our approach in the biologically more relevant task of predicting interactions for proteins that were not encountered during training.

## 2 Results

### 2.1 Proposed Approach

Most RPI prediction tools start by assigning a feature representation to proteins and RNAs, and then train a supervised machine learning algorithm on a set of annotated positive/negative interactions. While this has been a successful strategy by and large, it does not explicitly leverage the network information: protein/RNA nodes are simply summarised as feature vectors, and knowledge about shared targets/regulators is not directly incorporated in the prediction algorithm. With the advent of large CLIP-seq data sets, this network-level information is likely to play an increasingly relevant role in improving algorithmic performance. In this paper, we propose to adopt link prediction algorithms based on Graph Neural Networks (GNNs) to perform network-based prediction of RPIs. In recent years, GNNs have become an indispensable tool for applying neural networks to the graph domain (see Wu *et al*. (2020b) and Zhou *et al*. (2020) for recent reviews). GNNs use message passing between the nodes of a graph to non-linearly transform feature vectors and learn low dimensional embeddings for predicting interactions between nodes. In a way, GNNs generalize the well-known convolution operation of neural networks to graphs (Wu *et al*., 2020b; Zhou *et al*., 2020). In this paper, we use Graph Convolutional Network (GCN) (Kipf and Welling, 2017) with 2 convolutional layers as the GNN model (see Table S2 for a comparison with different GNN architectures).

Figure 1 illustrates the general workflow of our proposed framework. Briefly, CLIP-seq data for a specific cell line is used to identify RNA-protein interactions, thus creating an RPI network. Protein and RNA sequences are used to extract features for the nodes in the graph. RNA-seq for the cell line is used to identify abundant RNAs, which is subsequently used to identify highly likely negative interactions. The interactions and node features are then used to train a GNN (or other machine learning models), which transforms the features to learn low dimensional representations for the nodes and predicts interactions in different scenarios. In this paper, we use the Graph Convolutional Network (GCN) of Kipf and Welling (2017) as the encoder. Further details can be found in Section 4.

**Figure 1:**
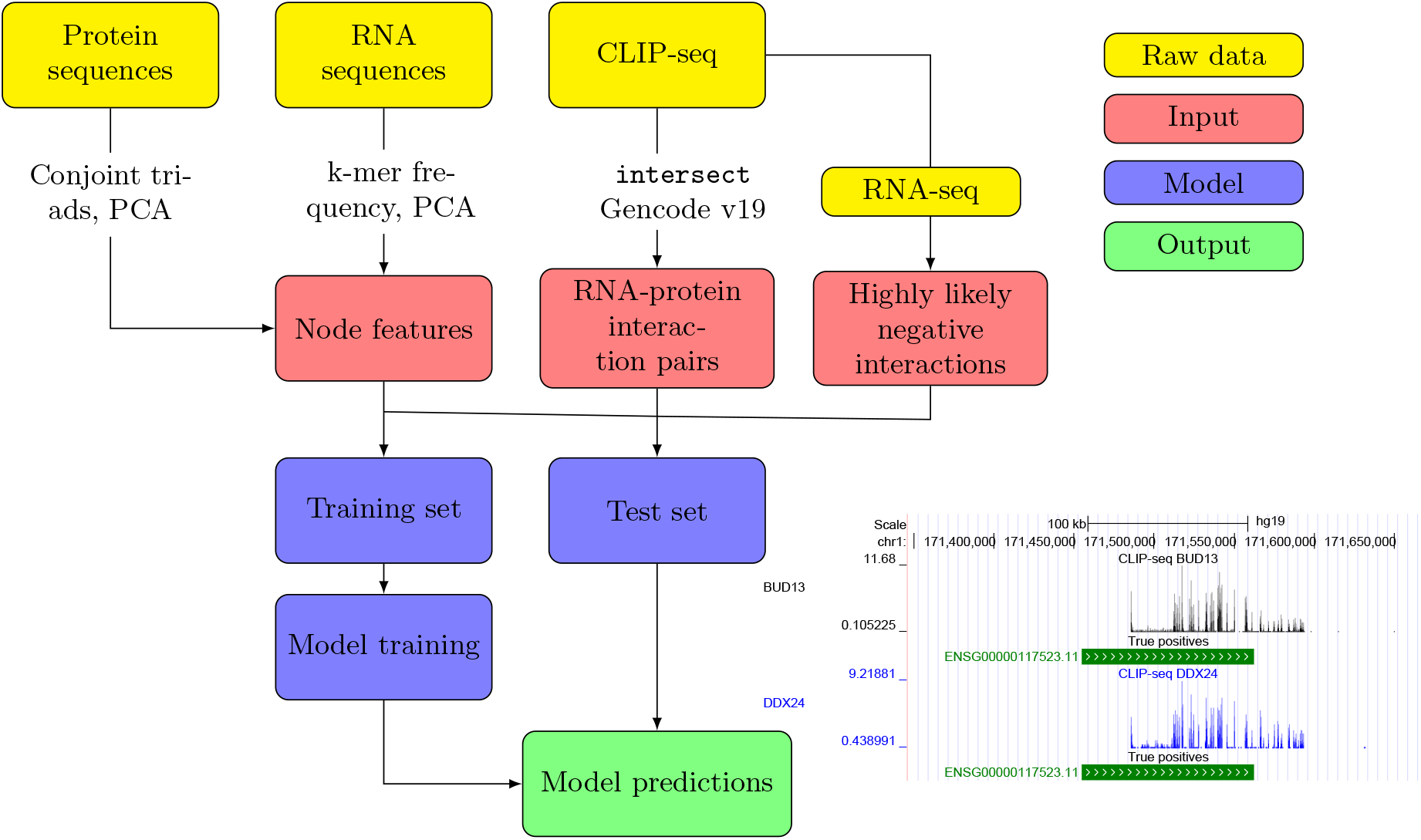
Pictorial representation of the framework presented in this paper: The raw data is transformed to obtain node features, positive and negative interactions, which serve as input for the GNN. The trained model is used for making predictions as shown using the genome tracks.

Our models were trained using the large-scale eCLIP datasets for RNA-protein interactions extracted from the ENCODE project (see Section 4.1). We use RPIseq (Muppirala *et al*., 2011) and RNAcommender (Corrado *et al*., 2016) as the baseline models to compare against our GNN-based approach. For GCN and RPIseq, we consider variants, GCN (RNA) and RPIseq (RNA), where RNA-seq information is used to provide information about the presence and quantity of RNA in a given biological sample (see Section 4 for more details). Other similar methods like IPMiner (Pan *et al*., 2016) and NPI-GNN (Shen *et al*., 2021) fail to run on our datasets due to memory and computation time issues.

The models are evaluated under three different scenarios: (i) we assume that some percentage of the RNA-protein interactions are missing in Section 2.2; (ii) we perform leave-one-protein-out experiments in Section 2.3, assuming the availability of full interaction information for the remaining proteins; and (iii) transfer learning of RNA-protein interactions from a source cell line to a target cell line in Section 2.4. In scenarios (ii) and (iii), we use the similarity between the sequence-based features of proteins to elicit the embedding for a previously unseen protein. Using machine learning terminology, we refer to scenario (i) as transductive learning (as the set on which predictions are needed is part of the graph), scenario (ii) as inductive learning, and scenario (iii) as transfer learning.

### 2.2 Transductive Link Prediction

For the first evaluation of the models, we consider the scenario when varying percentage of positive edges are removed from the RPI network. An equal number of negative interactions, as defined in Section 4.1.1, for the test set can be sampled either uniformly at random, or proportional to the degree of each protein. The second setting is considerably harder in practice because the network has to implicitly learn the degree information from training data. Additionally, the harder setting is likely to be more representative of true biological missing data. For GCN, we use the AUROC on the validation set to select the best model. We run all experiments 10 times and report the average results and standard deviations, highlighting in boldface the best results (determined using two sample t-tests) in tables.

A hyperparameter that needs to be tuned for GCNs is the dimension of the final embedding of the nodes. In Figure 2, we plot the performance of three different variants of GCN as the final embedding dimension is varied (the hidden dimension is kept constant at 50). While all methods for all embedding dimensions provide results which are clearly better than random predictors, the trend shows a clear peak at dimensions between 5 and 10 for all methods; we therefore choose 10 as the final embedding size for all subsequent evaluations of GCN under different settings. We also test the impact of increasing the number of layers from two to six in GCN. The results in Figure 2 show that the performance degrades as we stack more layers in GCN, which can be attributed to the over-smoothing of the node representations (Oono and Suzuki, 2019; Chen *et al*., 2020).

**Figure 2:**
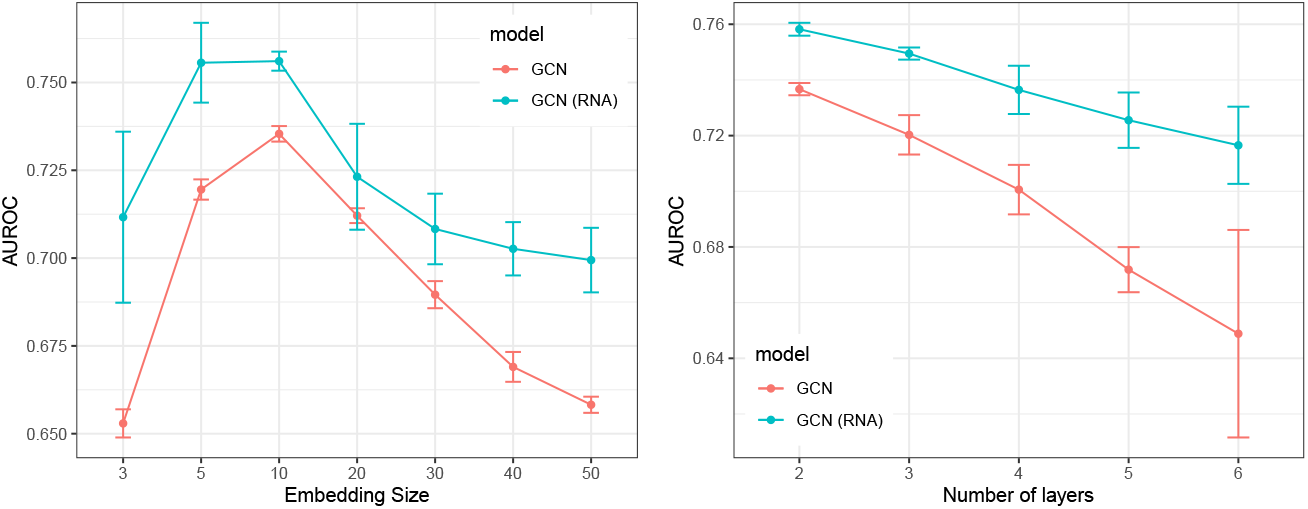
Comparing the performance of the two GCN settings while varying the size of the final embedding (left) and the number of layers in GCN (right). The test set contains 20% edges and the hidden embedding size is set to 50. The error bars show the standard deviation on 10 independent trials.

The results in Table 1 and 2 show the performance of different models on the transductive link prediction task for the ‘hard’ setting of negative interactions on the RPI networks of K562 and HepG2 cells, respectively. The tests are performed varying the percentage of edges in the test set from 10% to 50%. Our results show that our GCN-based models is comparable with RPISeq and consistently outperforms RNAcommender in the task of transductive link prediction by a clear margin. As expected, the performance drops as we increase the number of edges in the test set (thus decreasing the size of the training set), however for all sizes of training set the performance of the GCN approach remains above 70% AUROC. We observe that providing RNA-seq information consistently improves the predictive performance of both RPIseq and GCN (this effect is more pronounced when using word2vec based node features in GCN, as seen in Table S5).

**Table 1:** Comparing the AUROC for transductive learning setting in K562 cell line with varying percent of edges in the test set (validation set contains 10% edges in all cases). The bold marker denotes the best performing model(s) based on a t-test and ± denotes the standard deviation on 10 independent trials.

| Test | RNAcommender | RPISeq | RPISeq (RNA) | GCN | GCN (RNA) |
| --- | --- | --- | --- | --- | --- |
| 10% | 0.604±0.003 | **0.778**±0.003 | 0.774±0.004 | 0.751±0.002 | 0.768±0.004 |
| 20% | 0.601±0.004 | **0.767**±0.002 | **0.770**±0.004 | 0.736±0.002 | 0.757±0.002 |
| 30% | 0.595±0.004 | **0.759**±0.002 | **0.765**±0.003 | 0.717±0.002 | 0.740±0.002 |
| 40% | 0.594±0.005 | 0.749±0.002 | **0.761**±0.003 | 0.717±0.003 | 0.740±0.001 |
| 50% | 0.586±0.005 | 0.740±0.002 | **0.751**±0.003 | 0.709±0.003 | 0.730±0.002 |

**Table 2:** Comparing the AUROC for transductive learning setting in HepG2 cell line with varying percent of edges in the test set (validation set contains 10% edges in all cases). The bold marker denotes the best performing model(s) based on a t-test and ± denotes the standard deviation on 10 independent trials.

| Test | RNAcommender | RPIseq | RPIseq (RNA) | GCN | GCN (RNA) |
| --- | --- | --- | --- | --- | --- |
| 10% | 0.632±0.004 | **0.808**±0.003 | **0.805**±0.004 | 0.771±0.003 | 0.798±0.003 |
| 20% | 0.628±0.004 | **0.799**±0.002 | **0.800**±0.002 | 0.762±0.003 | 0.79±0.002 |
| 30% | 0.621±0.004 | **0.791**±0.001 | **0.796**±0.003 | 0.748±0.004 | 0.78±0.002 |
| 40% | 0.618±0.004 | 0.782±0.002 | **0.792**±0.004 | 0.742±0.004 | 0.772±0.002 |
| 50% | 0.618±0.004 | 0.773±0.001 | **0.784**±0.003 | 0.734±0.001 | 0.763±0.002 |

Results for the simpler setting when negative test edges are selected at random are shown in the Table 3. Here we see a significant improvement for all the approaches, with the GCN achieving test accuracies surpassing 90% (in some cases substantially so). In this case, the GCN outperforms RPIseq as well.

**Table 3:**
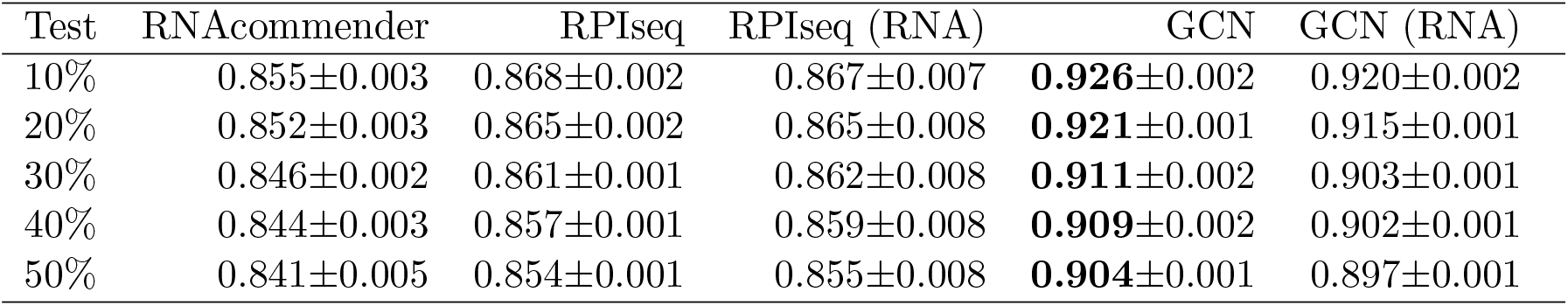
Comparing the AUROC for transductive learning setting in K562 cell line in ‘easy’ setting while varying the percent of edges in the test set (validation set contains 10% edges in all cases). The bold marker denotes the best performing model(s) based on a t-test. The error bar ± denotes the standard deviation of the test performance of 10 independent trials.

### 2.3 Inductive Link Prediction

The ability to make *de novo predictions* of RNA-protein interactions is one of the biggest motivations for developing computational models for this problem. This is known as the problem of inductive (or out-of-sample) link prediction in the GNN community. Such analysis is particularly valuable as the model predictions can serve as a proxy for proteins for which high-throughput data is not currently available. In this setting, we create a training network 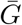 by removing interaction data for the test and validation proteins. The trained model then computes the embedding for a new protein v using normalized feature similarity sim(**x***_v_*, **x***_u_*) (based on inverse

Euclidean distance or cosine similarity) to previously seen proteins:

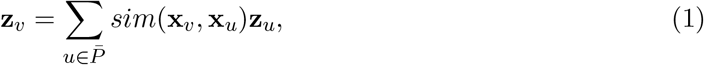

where 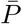 is the set of proteins in the training network 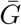, while **x***_u_* and **z***_u_* are the features and embeddings of a previously seen protein u. We perform experiments with a single protein in the test set (with all highly likely negative interactions as defined in Section 4.1.1). This can create a potential class imbalance in the positive and negative interactions in the test and validation sets. We therefore also use the average precision (AP) metric introduced in Section 4.4, which is a better measure for an imbalanced dataset. The protein with the highest feature similarity to the test protein is chosen as the validation protein. This is justifiable from a biological standpoint as it allows to reduce bias in the model predictions.

Figure 3 compares the performance of different models over the entire set of proteins in the inductive link prediction setting. Each box plots the distribution of mean AUROC/AP for proteins in the K562 cell line (10 replications for each protein). The results show that on average all variations of GCN outperform RNAcommender (labelled as RNAcom in the plots) in the K562 cell line. More specifically, GCN with inverse distance-based similarity outperforms RNAcommender on 93.33% and 87.5% of proteins on AUROC and AP, respectively. When compared with RPIseq, GCN with inverse distance-based similarity is better on 85% of proteins on both AUROC and AP. Among the different settings of GCN, we observe that the choice of similarity function has very little impact on the model performance and even appending RNA-seq to the final embeddings does not seem to substantially improve model performance (although we do see some improvement in the HepG2 cell line, see Figure S3).

**Figure 3:**
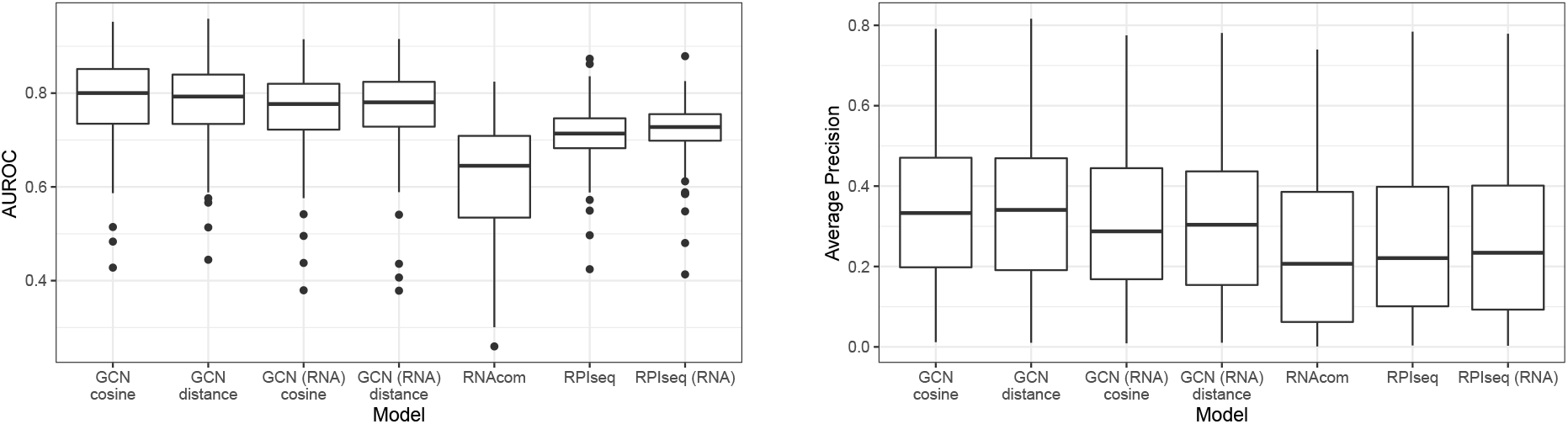
Comparing the performance of various models for de novo prediction in K562 cell line. Each box shows the distribution of mean AUROC (left) or average precision (right) over the entire set of proteins when the model is tested for a single protein in the test set.

Figure 4 shows genome tracks annotated by the predictions of our model for two proteins, BUD13 and DDX24, in the inductive link prediction task for the K562 cell line. Figures 4a and 4b show example regions corresponding to predicted true positives and true negatives; as expected, true positives correspond to regions with a strong binding signal, while true negatives display a complete absence of signal. Figure 4d and 4e show examples of wrong predictions (false positives and false negatives respectively): both examples show a modest amount of binding, likely representing regions that are borderline cases in the peak calling procedure. This suggests that the incorrect predictions by the model may correspond to potentially noisy regions. This point is illustrated globally using Figures 4c and 4f, which plot the distribution of reads per transcript for the four outcomes. We observe that the true positive and true negative predictions respectively have significantly higher and lower number of reads compared to the other cases, whereas the false positive and false negative predictions by the model have intermediate amounts of reads, potentially corresponding to noisy regions.

**Figure 4:**
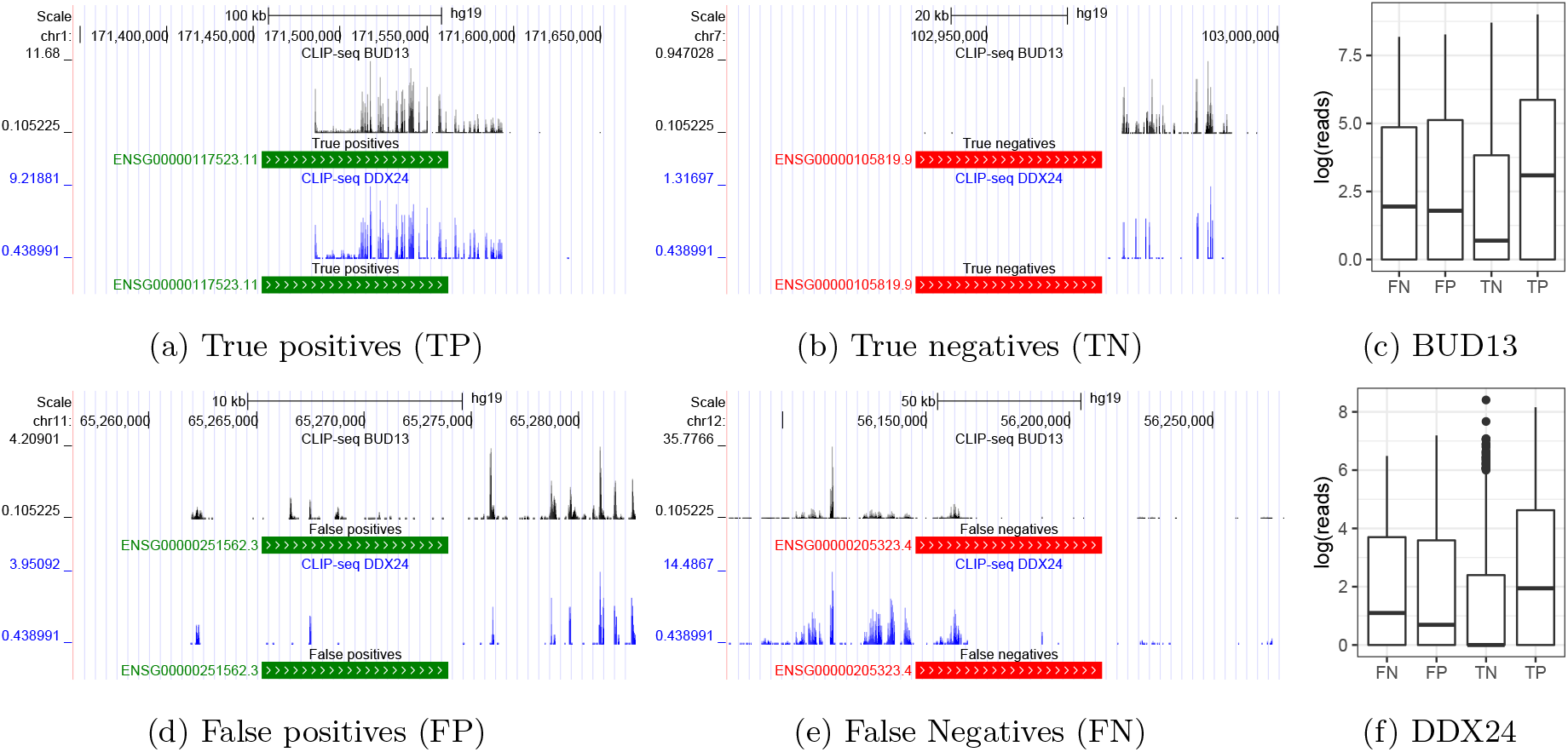
The plots show representative genome tracks produced using the eCLIP data annotated by predictions made by our model under four different outcomes. We consider two proteins, BUD13 and DDX24, in the inductive link prediction task for the K562 cell line. Positive predictions are shown in green and negative predictions in red. We also plot the distribution of reads (Figure 4c and 4f) for the two proteins under the four outcomes.

### 2.4 Transfer Learning

The ENCODE dataset (Van Nostrand *et al*., 2020) consists of 223 eCLIP experiments for 150 proteins across two different cell lines (K562 and HepG2). This provides an opportunity to perform transfer learning, where a model learnt from the eCLIP data for a source cell line can be used to predict the interactions in a target cell line. This is potentially the most interesting aspect of our approach, as it would permit researchers to obtain a reasonable prediction of an RPI network in new conditions based on minimal information about the target condition.

To train a model for transfer learning, we split the interactions from the source cell line by assigning all interactions from a fixed percentage of randomly chosen proteins to the validation set. Splitting the data in this way allows us to choose a model that has higher predictive power on previously unseen proteins. For creating the test set using the target cell line, we only consider interactions with RNAs that already exist in the source cell line. This allows us to exclusively focus on transfer learning for proteins. Negative interactions in the test set (same number as positive interactions) are sampled uniformly at random from the highly likely negative interactions in the target cell line (‘easy’ setting in Section 2.2). This is a reasonable assumption because to sample negative interactions proportional to a protein’s degree (‘hard’ setting), we need to have a priori knowledge about its interactions in the target cell line.

As in Section 2.3, we use Equation 1 to compute embeddings for the proteins in the test and validation sets. We specifically focus on the use of RNA-seq in transfer learning as it can provide information about RNA abundance in the target cell line. For transfer learning, we append the transcript per million (TPM) counts of the source cell line to the RNA embeddings during training, but replace it with the TPM counts in the target cell line for making the final predictions. Note that RNAcommender cannot be used for transfer learning^1^, which is why it has been omitted in Tables 4 and 5.

**Table 4:** Comparing the AUROC for transfer learning from K562 to HepG2. We vary the percentage of proteins in the validation set of the source cell line, thus reducing the number of interactions (and proteins) in the training network. The bold marker denotes the best performing model(s) based on a t-test and ± denotes the standard deviation on 10 independent trials.

| % val | RPIseq | RPIseq (RNA) | GCN | GCN (RNA) |
| --- | --- | --- | --- | --- |
| 5 | 0.697±0.003 | 0.689±0.003 | 0.693±0.024 | **0.724**±0.009 |
| 10 | 0.697±0.004 | 0.688±0.004 | 0.688±0.026 | **0.719**±0.006 |
| 20 | **0.696**±0.004 | 0.686±0.004 | 0.677±0.022 | **0.704**±0.008 |
| 30 | **0.695**±0.006 | **0.688**±0.004 | 0.664±0.016 | **0.689**±0.011 |

**Table 5:** Comparing the AUROC for transfer learning from HepG2 to K562. We vary the percentage of proteins in the validation set of the source cell line, thus reducing the number of interactions (and proteins) in the training network. The bold marker denotes the best performing model(s) based on a t-test and ± denotes the standard deviation on 10 independent trials.

| % val | RPIseq | RPIseq (RNA) | GCN | GCN (RNA) |
| --- | --- | --- | --- | --- |
| 5 | 0.706±0.005 | 0.614±0.011 | **0.785±0.037** | **0.789±0.017** |
| 10 | 0.705±0.003 | 0.616±0.007 | **0.785±0.026** | **0.778±0.016** |
| 20 | 0.705±0.004 | 0.617±0.010 | **0.774±0.018** | **0.760±0.018** |
| 30 | 0.704±0.004 | 0.616±0.009 | **0.759±0.021** | **0.748±0.016** |

The results in Tables 4 and 5 show that the GCN approach provides a good predictive accuracy even in the transfer learning mode, with AUROC values over 70% in most of the cases. Further, the best GCN variant also outperforms RPIseq in the transfer learning scenario. It should be noted that these results should be compared to Table 3 as we use the ‘easy’ setting for negative edges in this section. Additionally, there appears to be a clear advantage of using RNA abundance data in transfer learning as GCN (RNA) is the best performing model in most cases. This is intuitively appealing, as it shows that the RNA-seq data clearly conveys some information about the state of the cell which is relevant to the prediction of the RPI network. Nevertheless, it is still very surprising that, even without RNA-seq information, GCN provides good predictive performance. To contextualise this observation, in Figure 5 we compare the ROC curve for GCN (RNA) to the prediction we would obtain by just assuming the two RPI networks to be the same on the set of proteins/RNAs shared by the two eCLIP experiments (naive transfer). Remarkably, the performance of this naive transfer approach is only marginally better than random, and considerably worse than the GCN prediction at the same false positive rate. This indicates that the GCN learns primarily *robust* interactions that are seen in multiple different environments, which are presumably hard-wired into the protein-RNA sequence features.

**Figure 5:**
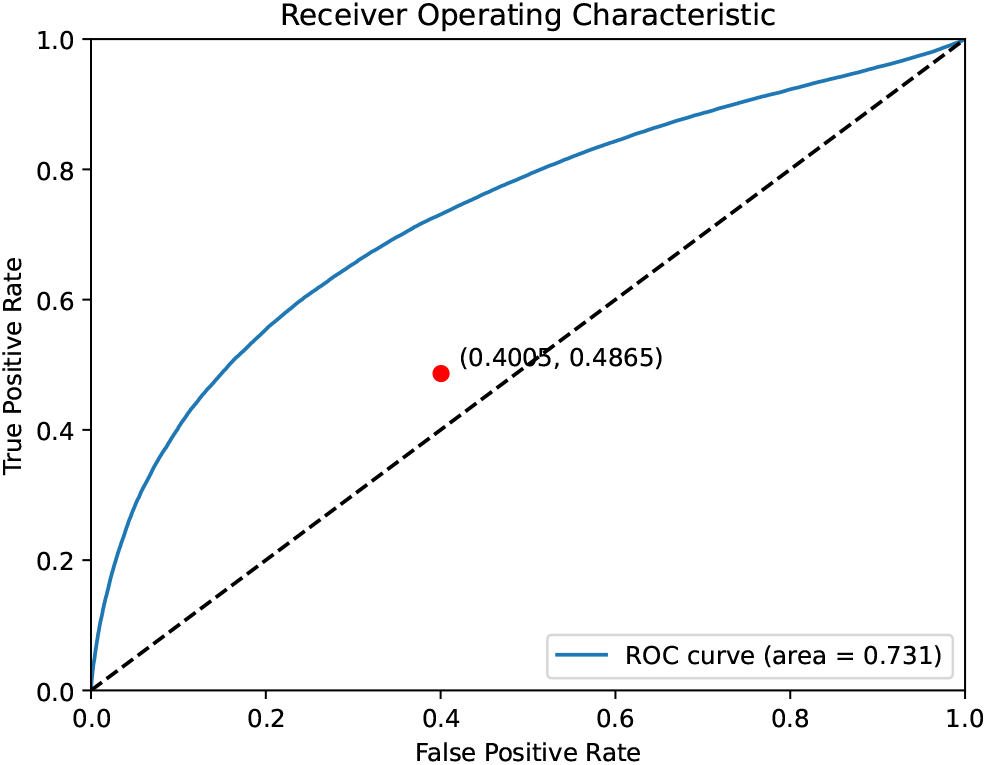
ROC curve for transfer learning from K562 to HepG2 cell line for GCN (RNA). Red dot corresponds to the false positive and true positive rates if the edges from the source cell line are directly transferred to target cell line.

We also observe that the performance of GCN degrades as the size of the validation set is increased. This implies that the model learns better by seeing more data from the source cell line instead of just overfitting to the training data. Comparing the results in Table 4 and 5, we observe that transfer from HepG2 to K562 seems easier than in the other direction, specifically for GCN. There are obvious differences between the two datasets: more proteins were profiled within the K562, while the HepG2 dataset had higher density of interactions, both of which may affect the complexity of the models learnt. Additionally, the two biological systems are very different (one a leukaemia cell line, the other derived from a hepatocellular carcinoma), likely with different complexities. None of these observations however points to a clear reason why transfer should work better in a direction than the other. A follow-up study trying to investigate these modest difference in performance would likely need a deeper dive in the biology and novel experiments, and is thus outside the scope of this paper.

## 3 Discussion

Experimental discovery of RPIs has been a major focus of research over the last ten years. After a pioneering period where novel technologies were still being developed, the last few years have seen an effort towards scaling and standardising the technology (Van Nostrand *et al*., 2016), resulting in the publication of the first large-scale compendia of RNA-RBP interactions in human cell lines (Van Nostrand *et al*., 2020).

These technological developments call for a change in the way RPI data are modelled. Earlier approaches (reviewed in e.g. (Pan *et al*., 2019)) focused on predicting the targets (or the binding sites) of individual RBPs, treating potential target transcripts as i.i.d. observations thus enabling the deployment of machine learning approaches such as GNNs. Even when a network of interactions was used (for example, in RPIseq Muppirala et al. (2011)), the datasets did not consider genome-wide targets for proteins and were hence incomplete. Now, the availability of binding data from hundreds of RBPs leads to hundreds of correlated prediction tasks, calling for methods that can effectively leverage the network of interactions. In this spirit, our GNN model transfers information between different RBPs binding data, translating the problem of predicting the binding targets of an RBP to predicting the whole RPI network.

Our experiments demonstrate considerable promise in this attempt. While our GNN-based approach is on-par or better than other competitors on the classical tasks of link prediction, it offers strong predictive performance in out-of-sample inductive predictions of the targets of an unseen RBP. Additionally, we show empirically that the GNN approach is also able to perform transfer learning, i.e. predict the RPI network in a different (related) condition starting from RPI data from a well characterised condition, a task that was never attempted to our knowledge. Further, we found that the use of RNA-seq information enhanced the performance of the GNN model. Overall, our approach provides significant advantages and should be chosen for making *de novo* predictions of RNA-protein interactions, specifically when bolstered by the use of RNA-seq information.

While we believe GNNs hold much promise for the problem of RPI network prediction, a number of areas for future improvement are clearly open. First of all, proteins (and transcripts) are characterised solely by their sequence and their binding partners in our approach, making the task of predicting the full complement of binding partners for a new protein (inductive link prediction) difficult. In principle, the availability of additional node information (for example in the form of protein-protein interactions, or of ontological annotations) could be easily incorporated in the GNN framework, potentially leading to significant improvements. Another area of great interest where improvements are certainly possible is transfer learning. Here the question is to identify suitable covariates which can be used to measure the similarity of different domains. In this paper, we show that the use of RNA-seq data helps in the transfer learning task, presumably because it recapitulates some information on the state of the cell, nevertheless more appropriate task descriptors might be found and integrated in the framework. While we found that a simple GNN architecture can provide strong predictive performance, there remains a need to further investigate heterogeneous GNN models and/or architectures specifically designed for link prediction. Finally, using richer features derived from structural information of proteins and RNAs is another aspect that can potentially improve performance of machine learning methods.

## 4 Materials and Methods

### 4.1 Dataset

CLIP-seq experiments can provide genome-wide binding sites for RBPs. To retrieve these binding sites, the CLIP-seq reads are first mapped to a reference genome, followed by identification of the peaks of reads using peak calling softwares. These peaks correspond to RBP binding sites based on a certain predefined cutoff, which can be used to identify the set of RNAs a protein binds to.

To construct the benchmark datasets, we use the eCLIP dataset for two cell lines (HepG2 and K562) generated as part of the ENCODE project phase III (Van Nostrand *et al*., 2020). We use the highly reproducible peaks identified from the two replicates of the eCLIP data using the Irreproducibility Discovery Rate (IDR) framework (Li *et al*., 2011) to obtain the binding regions. The gene corresponding to the binding site is obtained by using the bedtools intersect function (with a pre-defined minimum overlap between genomic features, we use 50% in this study) with the human genome (Gencode v19 is used). Repeating the procedure for each protein in the eCLIP dataset we obtain a bipartite network *G* = (*V, E*, **X**) of RNA-protein interactions for a particular cell line. The network G contains a set of nodes *V* = *R* ∪ *P*, where *R* is the set of RNAs and P is the set of RBPs, an edge set *E* ⊆ *R* × *P* of RNA-protein interactions, and a matrix **X** of node features (see Section 4.1.2 for further details).

The final graph for the K562 cell line consists of 14665 nodes (120 proteins and 14545 RNAs) with 144527 interactions between proteins and RNAs. The mean (out) degree of proteins is 1204.39 with a standard deviation of 1304.64, while the mean and standard deviation of the RNA (in) degree are 9.94 and 10.27, respectively. For the HepG2 cell line, the graph contains 15018 nodes (103 proteins and 14915 RNAs) and 145509 edges. The mean (out) degree of proteins is 1412.71 with a standard deviation of 1380.69, while the mean and standard deviation of the RNA (in) degree are 9.76 and 10.03, respectively. Figure S1 plots the distribution of protein and RNA degrees for the RPI networks in the two cell lines. As far as we know, we are the first ones to create a graph from a homogeneous dataset with hundreds of RBPs using the eCLIP dataset of Van Nostrand *et al*. (2020).

#### 4.1.1 Negative Interactions

CLIP-seq experiments provide information about binding sites from the peaks of reads, but they do not provide any information about unbound sites. False negatives in a well-known problem in CLIP-seq data because of absence or low concentration of transcripts in the cell line used for experiments (Uhl *et al*., 2017; Maticzka *et al*., 2014). Appropriately defining negative samples is important for training machine learning algorithms (Mikolov *et al*., 2013a). Negative interactions play a crucial role in computation of the loss function, and recent work by Ying *et al*. (2018) has shown that appropriately choosing the negative samples can boost the performance of a link predictor. This becomes even more important for bipartite networks where randomly sampling two unconnected nodes can produce edge-types that do not exist in the data (RNA-RNA for example).

To tackle this issue, the following two strategies have been commonly used to construct negative samples from CLIP-seq data (Pan *et al*., 2019): (i) use the regions where no binding sites are located as negative samples, or (ii) use randomly shuffled nucleotides in the positive sequences as negative samples. We augment the first strategy by utilizing the RNA-seq transcript abundance data to identify RNAs that have transcripts per million (TPM) counts more than the median value in the cell line but still do not have any peaks with the corresponding protein to define negative samples. This strategy allows the model to learn from highly likely unbound sites of real RNA sequences and alleviates the problem of false negatives described above. Using RNA-seq we identify reliable non-interacting RNAs for each protein and consequently use these negative samples to define the training and test sets.

#### 4.1.2 Node Features

Node features are essential for training GNNs as they allow the neighborhood aggregation process to capture the hierarchical non-linearities in the network data. The sequences of proteins and RNAs can be encoded as numeric vectors for training machine learning models. There are two common ways of encoding RNA sequences as numeric vectors are (Pan *et al*., 2019): (i) one-hot vector, which is a bit vector that consists of all zeros except for a single dimension, and (ii) k-mer frequency vector, which is a vector consisting of frequencies of all k-mers, similar to bag-of-words (BOW) in natural language processing. We use the following feature extraction methods, which have been successfully used by previous methods for predicting RNA-protein interactions (Muppirala *et al*., 2011; Pan *et al*., 2016):

- **Proteins**: Conjoint triad descriptors (Shen *et al*., 2007) abstract the features of proteins based on the classification of amino acids according to their dipoles and volumes of the side chains. Each protein sequence is encoded using a normalized 3-gram frequency distribution extracted from a 7-letter reduced alphabet representation.
- **RNA**: k -mer frequency distribution counts the frequency of individual k-mers (AAA, AAC, …, UUU are 3-mers) in a given RNA sequence. It is the simplest feature extraction method for RNAs, where *k* can be used as a hyperparameter. We use *k* = 6 for our experiments.

After obtaining the protein and RNA sequences for the nodes in the network (see Section S1 for more details), the aforementioned feature extraction methods are used to obtain 7^3^ = 343 and 4^6^ = 4096 dimensional feature vectors for proteins and RNAs, respectively. For aggregating the features in a homogeneous GNN model such as GCN, node features should have the same dimensionality. To achieve this, we use the first 100 principal components of protein and RNA features to define the node features **X** for the network *G*. To account for the heterogeneous nature of the nodes in the network (proteins and RNAs), we append 1 and 0 to the protein and RNA features, respectively. This creates a separation in the feature space of proteins and RNAs, thus allowing the GNN to distinguish between the two types of nodes.

Features can also be extracted by treating RNA and protein sequences as a special kind of language, where k-mers can be treated as words and sequences as sentences. Natural language processing techniques such as word2vec (Mikolov *et al*., 2013b) can then be used to learn embeddings for protein and RNA sequences (Asgari and Mofrad, 2015). Results for this alternative feature extraction scenario can be found in the Supplementary information.

### 4.2 Baselines

We use RPIseq (Muppirala *et al*., 2011) and RNAcommender (Corrado *et al*., 2016) as the baseline models to compare against our GNN-based approach. RNAcommender is a recommender system capable of suggesting genome-wide RNA targets for unexplored RBPs using interaction information available from high-throughput experiments performed on other proteins. In our evaluations, we use sequence-based features instead of the advanced feature engineering^2^ (protein domain composition and the RNA predicted secondary structure) used in the original implementation of RNAcommender (Corrado *et al*., 2016). The sequence-based features that are easier to collect and also ensure that all the methods compared in this paper use the same input features.

RPIseq (Muppirala *et al*., 2011) uses a family of classifiers to predict RNA-protein interactions using only sequence information. RPIseq uses 4-mer frequency distribution counts in a given RNA sequence as RNA features, while Conjoint triad descriptors (Shen *et al*., 2007) are used for creating protein features. In our experiments, we use a random forest with 20 trees, which was shown to outperform an SVM-based classifier in the original publication (Muppirala *et al*., 2011). For RPIseq, we consider a variant, RPIseq (RNA), where RNA-seq information is appended to the features in order to provide information about the presence and quantity of RNA in a given biological sample. This allows us to perform a fair comparison when both RPIseq and GCN are supplied with the additional RNA-seq information.

### 4.3 Graph Neural Networks

The ability to learn from the entire network of RNA-protein interactions enables us to build a single end-to-end model for predicting RNA-protein interactions. The GNN architecture creates a non-linear permutation invariant transformation function on node, edge and graph features, which can be optimized for performing downstream learning tasks. The neighborhood aggregation process of GNNs allows us to capture the hierarchical non-linearities in network data, thus learning low dimensional embeddings for the nodes of a graph. Further, the GNN architecture facilitates the aggregation of information from distant neighbors such as other proteins, thus learning better node representations. A lot of the existing GNN architectures can be directly translated into the framework of message passing neural networks (Gilmer *et al*., 2017), where each node sends and receives messages (using function *M_k_*(·)) from its neighbors, and subsequently updates (using function *U_k_*(·)) its own node state:

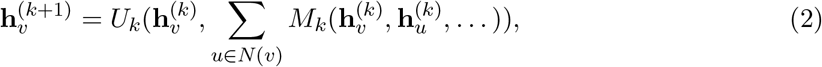

where 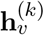 is the hidden representation of node *v* in layer *k, M_k_*(·) and *U_k_*(·) are functions with learnable parameters, *N*(*v*) is the set of neighbors of node v, and … represent other features (such as edge features) that can be added to the message passing process. Node features, if available, can be used as the initial hidden representation 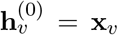 for a node. After *K* message passing layers, node embedding **z***_v_* is produced for each node *v* as the final output, which can then be used for node, link, or graph level prediction tasks. The functions *M_k_*(·) and *U_k_*(·) share parameters across nodes, but each node is associated with an individual computation graph defined by its neighbors (Ying *et al*., 2019).

In this paper, we use Graph Convolutional Network (GCN) (Kipf and Welling, 2017) with 2 convolutional layers as the GNN model. GCN bridges the gap between spectral and spatial approaches for performing convolution over graph-based data (Wu *et al*., 2020b). The graph convolution operation of GCN can be written as:

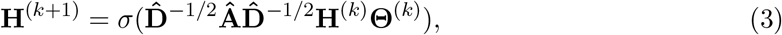

where 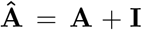 is the adjacency matrix with self loops and 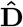 is the diagonal degree matrix corresponding to 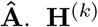 is the matrix containing the hidden representation of nodes at layer *k* with 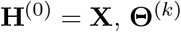 are the model parameters at layer *k*, and *σ*(·) is an element-wise activation function.Comparing Equations 2 and 3, for GCN the message and update functions take the following form (Gilmer *et al*., 2017):

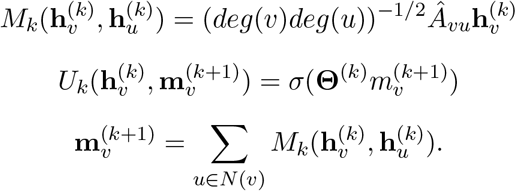

Shchur *et al*. (2018) performed a comprehensive analysis of different GNN architectures and found that there is no clear winner when it comes to choosing a GNN architecture, at least on the benchmark datasets. Our results in Table S2 also show that there is little difference between the performance of various GNN architectures. Following these observations, we decided to choose the simplest architecture with the fewest parameters for our experiments.

To add some biological context to the model, we consider using RNA-seq information for GCN. RNA-seq experiments provide high resolution information about the presence and quantity of all the RNAs in a given biological sample. RNA-seq can tell us which genes are turned on in a cell, and what their level of transcription is (Ozsolak and Milos, 2011). Thus, one would assume that RNA abundance would be a good indicator of an RNA’s ability of being bound by a protein. In GCN (RNA), we append log(1 + TPM) to the final node embedding **Z** of RNAs (set to −1 for all proteins) obtained after GCN layers. The modified embeddings are then used for computing the loss and predicting interactions.

### 4.4 Link Prediction and Evaluation Metrics

The current knowledge of interactions in biological networks is often incomplete, which makes predicting missing interactions an important task (Muzio *et al*., 2020). Given the importance of RNA-protein interactions and the challenges associated with experimental methods, predicting potential interactions using computational models can compliment our current knowledge (Corrado *et al*., 2016). While most existing studies focus on transductive link prediction (both nodes are in the graph), inductive (or out-of-sample) link prediction can prove immensely valuable for new proteins. Link prediction is often framed as a semi-supervised learning problem, where the known links in a network are used to predict additional interactions (Al Hasan *et al*., 2006; Liben-Nowell and Kleinberg, 2007). With the rise in the popularity of GNNs, specialized methods (Kipf and Welling, 2016; Zhang and Chen, 2018; Zhang *et al*., 2020) have been developed to deal with the link prediction task using GNNs. The link prediction task introduces additional challenges for GNNs as the method needs to learn link representations instead of node representations (Zhang *et al*., 2020). Link prediction is an important task for recommender systems (Wu *et al*., 2020a) and has been well-studied in the heterogeneous graph learning literature (Yang *et al*., 2020; Wang *et al*., 2020).

In a GNN for link prediction, the message passing procedure described in Section 4.3 is used to compute individual node representations **z***_u_*, following which a function *p_uv_* = *f* (**z***_u_*, **z***_v_*) can be used to predict the probability of the link (*u, v*). In our implementation, we use the dot product of the final embeddings as the function *f* (·) The model can be trained to maximize the likelihood of reconstructing the true adjacency matrix **A** using the binary cross entropy loss:

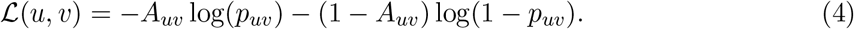

Splitting networks into training and test subnetworks is not trivial in link prediction problems. While performing the train-test split of edges, we need to make sure that every node in the training network has a non-empty set of neighbors so that the GNN can learn appropriate representations using the message passing process shown in Equation 2. To ensure this, test edges are sampled for each RNA while making sure that it stays connected in the training network.

Link prediction is a binary classification task and the performance of an algorithm can be evaluated using different metrics. These metrics can be divided into two broad categories: fixed-threshold metrics and threshold curves (Yang *et al*., 2015). In research context, we generally do not have a reasonable threshold, which is why threshold curves and scalar measures summarizing them are widely used in the literature (Davis and Goadrich, 2006; Clauset *et al*., 2008; Lichtenwalter *et al*., 2010). In this paper, we use area under the receiver operating characteristic (AUROC), and average precision (AP) to evaluate performance of different methods on the link prediction task. The receiver operating characteristic (ROC) curve represents the performance trade-off between true positives and false positives at different decision boundary thresholds. AUROC reflects the probability that a randomly chosen positive instance appears above a randomly chosen negative instance. AP summarises the precision-recall curve, and is a better measure for a highly imbalanced dataset (Davis and Goadrich, 2006; Yang *et al*., 2015). AP can be computed using the following formula:

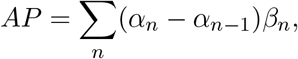

where *β_n_* and *α_n_* are the precision and recall at the *n^th^* threshold.

## Acknowledgements

The authors want to thank Gianluca Corrado for sharing the implementation details of RNAcommender.

## Footnotes

1 this conclusion was reached after discussions with the authors of RNAcommender.

2 this was done because it was not possible to perform the feature engineering for all the RNAs and proteins in our new dataset.

