## Supplementary Information for "*De novo* prediction of RNA-protein interactions with Graph Neural Networks"

Viplove Arora and Guido Sanguinetti

July 15, 2022

#### S1 Data Processing

1. Download the following data:
   - eCLIP .bed files using the accession identifier ENCSR456FVU at the ENCODE Data Coordination Center (`https://www.encodeproject.org`).
   - Comprehensive gene annotations for Human `Gencode v19` and the genome sequence (GRCh37.p13) from `https://www.gencodegenes.org/human/release_19.html`.
   - Obtain protein sequences from `UniProt` (`https://www.uniprot.org/proteomes/UP000005640`).
   - RNA-seq for K562 (`https://www.encodeproject.org/experiments/ENCSR109IQO/`) and HepG2 (`https://www.encodeproject.org/experiments/ENCSR181ZGR/`) cell lines.
2. Convert the Human Genome to .bed format using `gtf2bed` and extract only the gene regions from the genome. Extract the sequence of the genes from the entire genome sequence.
3. For each eCLIP experiment, use `bedtools intersect` on the highly reproducible peaks identified from the two replicates of the eCLIP data using the Irreproducibility Discovery Rate (IDR) framework (Li *et al.*, 2011) to find intersections with extracted gene regions and thus identify RNA-protein interactions.
4. Use the RNA-protein interactions to create separate bipartite networks for K562 and HepG2 cell lines.
5. Use the protein and RNA sequences to create node features as described in Section 4.1.2.

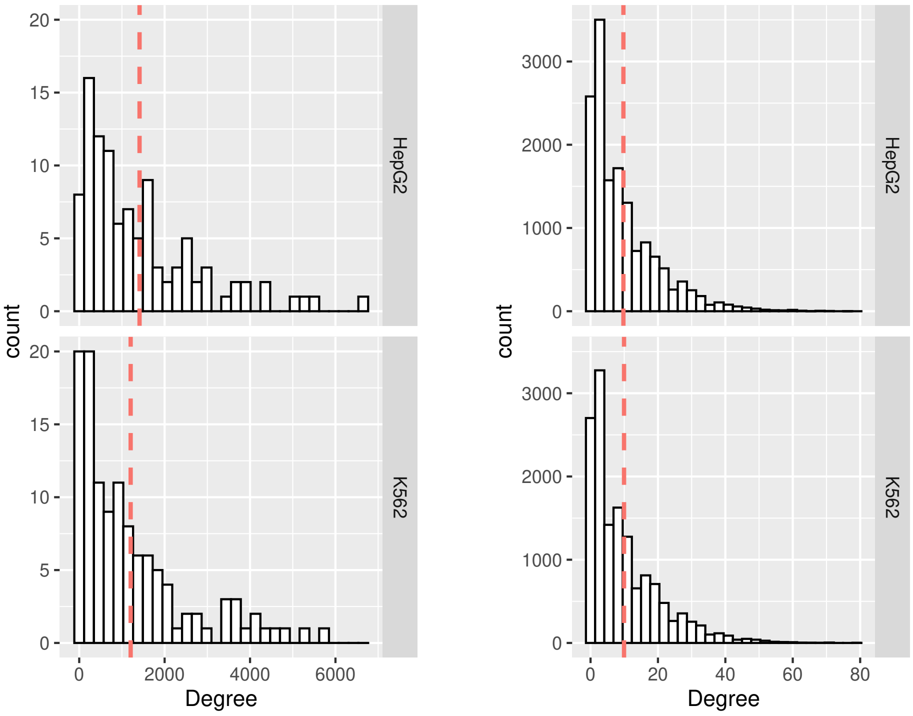


Figure S1: Distribution of protein (left) and RNA (right) degrees for the RPI networks in the two cell lines. The dashed red line shows the mean degree.

As far as we can tell, all models for RNA-protein interaction prediction, such as RPIseq (Muppirala *et al.*, 2011) and IPminer (Pan *et al.*, 2016) have been designed for small datasets with a handful of proteins and targets (see Table S1). This is easily explained by the fact that until very recently large, homogeneous datasets with hundreds of RBPs were simply not available; nevertheless, we found that the architectures and models employed so far had very poor scalability. Even recent methods like NPI-GNN (Shen *et al.*, 2021) are being designed to deal with these small datasets and run into computational issues when tested on bigger datasets like ours. The only method we found that was developed with large datasets in mind was RNAcommender (Corrado *et al.*, 2016) (which in the original paper was trained and tested on heterogeneous data from different experiments). Hence our choice of RNAcommender as the baseline model. Additionally, Corrado *et al.* (2016) also showed that their model outperforms previous models on larger datasets.

The network of interactions between RNAs and proteins forms a bipartite graph (see Figure S1 for distribution of node degrees). In the GNN literature, this falls in the category of heterogeneous graph, which has been the topic of some recent surveys on heterogeneous graph representation learning (Yang *et al.*, 2020; Wang *et al.*, 2020).

**Definition 1 (Heterogeneous graph)** *A heterogeneous graph is defined as a graph* $G = \{V, E\}$*, in which*

Table S1: Size of different datasets used for RNA-protein interaction prediction. Note that RNAcommender predicts interactions between proteins and UTRs instead of RNAs.

| Paper | Dataset | Interactions | Transcripts/UTRs | Proteins |
|---|---|---|---|---|
| NPI-GNN | NPInter2.0 | 10412 | 4636 | 449 |
| | RPI7317 | 7317 | 1874 | 118 |
| | RPI2241 | 2241 | 838 | 2040 |
| | RPI369 | 369 | 331 | 338 |
| IPMiner | NPInter2.0 | 10412 | 4636 | 449 |
| | RPI13254 | 13254 | 4500 | 42 |
| | RPI1807 | 1807 | 1078 | 1807 |
| | RPI2241 | 2241 | 842 | 2043 |
| | RPI488 | 243 | 25 | 247 |
| | RPI369 | 369 | 332 | 338 |
| RNAcommender | AURA2 | 502178 | 72226 | 67 |
| Ours | K562 | 144527 | 14545 | 120 |
| | HepG2 | 145509 | 14915 | 103 |

*$V$ and $E$ represent the node set and the link set, respectively. Each node $v \in V$ and each link $e \in E$ are associated with their mapping function $\psi(v) : V \rightarrow B$ and $\psi(e) : E \rightarrow C$. $B$ and $C$ denote the node types and link types, respectively, where $|B| + |C| > 2$.*

##### S1.1 Edge Weights

The procedures described above creates a binary RNA-protein network, but a protein can potentially have multiple CLIP-seq reads with an individual RNA. The multiple reads could either be a result of high concentration of the RNA transcripts or due to higher affinity towards the target RNA. To incorporate this information in our RPI networks, we define edge weights as a function of the number of observed interactions (CLIP-seq reads) normalized by RNA abundance obtained from RNA-seq TPM counts. For each protein, we use the empirical cumulative distribution function of the ratio between the number of reads and TPM count for the interacting RNAs to define the edge weights. These edge weights can then be utilized to create a weighted adjacency matrix instead of a binarized one in Equation 3. This will allow the model to prioritize information propagation along edges with higher weight.

#### S2 Methods

##### S2.1 Other GNN architectures

We consider three other GNN-based models in addition to GCN described in Section 4.3. Two of these architectures, R-GCN and HGT, are specifically designed for performing prediction tasks in heterogeneous graphs.

- **VGAE:** Graph auto-encoders (Kipf and Welling, 2016) use an encoder to learn low-dimensional vector representation of nodes and use a decoder to reconstruct the adjacency matrix from the latent representations. Variational GAE instead learns the distribution of data to avoid overfitting.
- **GIN:** Graph isomorphism network (Xu *et al.*, 2018) uses multi-layer perceptrons to create an injective function for neighborhood aggregation, thus creating a GNN with maximum discriminative power.
- **R-GCN:** Relational graph convolutional network (Schlichtkrull *et al.*, 2018) uses different convolutional layers for each node and edge type to update node representation by aggregating the representation vector of neighboring nodes through a normalized sum.
- **HGT:** In the heterogeneous graph transformer, (Hu *et al.*, 2020) propose to use the edge type to parameterize a Transformer-like self-attention architecture. The self-attention mechanism is accompanied by a message passing process (as described in Section 4.3) while also incorporating node and edge types.

GCN, GIN, and VGAE implementations are taken from PyTorch Geometric[1] (Fey and Lenssen, 2019) python library, while for R-GCN and HGT we use the Heterogeneous Network Representation Learning benchmark code[2] provided as a part of the recent survey (Yang *et al.*, 2020).

#### S2.2 Structured Negative Sampling

As described in Section 4.1.1, negative sampling plays an important role in the training of machine learning models. In the default training regime for GCN, negative edges are sampled uniformly at random from the set of possible negative interactions, which could result in a case where more negative interactions are sampled for a protein with a low degree. To deal with this issue, we consider structured negative (SN) sampling, where negative interactions for a protein are sampled proportional to its degree.

#### S2.3 Ranking-based loss

In addition to the GCN settings described in Section 4.3, we also consider an alternative loss function. The ranking (or contrastive) loss function aims to predict relative distances between inputs. In link prediction, this corresponds to comparing a positive interaction to a negative interaction. Ranking-based loss $\mathcal{L}_R$ can be computed using the following formula:

$$\mathcal{L}_R(u, v) = A_{uv}||\mathbf{z}_u - \mathbf{z}_v|| + (1 - A_{uv}) \max(0, m - ||\mathbf{z}_u - \mathbf{z}_v||), \tag{S1}$$

[1]https://github.com/pyg-team/pytorch_geometric
[2]https://github.com/yangji9181/HNE

where $m$ is the minimum distance required for negative interactions. In our experiments we compute the loss between a positive interaction for a protein with a corresponding negative one. Because of this, ranking loss can only be appropriately used with structured negative sampling.

### S3 Additional Results

We first compare the results for different GNN architectures described in Section S2.1 in the transductive learning setting. The results in Table S2 surprisingly show that homogeneous GNN architectures such as GCN, GIN, and VGAE perform significantly better than those designed for heterogeneous graphs.

Table S2: Comparing the AUROC for transductive learning setting in K562 cell line for different GNN architectures while varying the percent of edges in the test set (validation set contains 10% edges in all cases). The bold marker denotes the best performing model(s) based on a t-test. The error bar ± denotes the standard deviation of the test performance of 10 independent trials.

| Test | GCN | GIN | HGT | R-GCN | VGAE |
|---|---|---|---|---|---|
| 10% | **0.747**±0.005 | **0.743**±0.003 | 0.610±0.003 | 0.614±0.006 | **0.742**±0.005 |
| 20% | **0.735**±0.003 | **0.732**±0.002 | 0.598±0.001 | 0.610±0.008 | **0.729**±0.004 |
| 30% | **0.716**±0.003 | **0.717**±0.003 | 0.589±0.014 | 0.610±0.006 | 0.708±0.004 |
| 40% | **0.718**±0.001 | **0.717**±0.002 | 0.594±0.002 | 0.612±0.005 | 0.709±0.003 |
| 50% | **0.708**±0.001 | **0.706**±0.002 | 0.602±0.001 | 0.611±0.004 | 0.696±0.004 |

Following the results in Table S2, we decided to choose GCN as our primary model. We consider the following settings for GCN: vanilla version (GCN), weighted edges (W), RNA-seq appended to final node embeddings (RNA), structured negative sampling during training (SN), and ranking-based loss function (RL), along with all possible combinations of these settings. This results in 12 different settings for GCN (because RL can only be used with SN, as previously described in Section S2.3), as shown in Tables S3 and S4. A subset of these results (for the best performing GCN settings) can also be in the main text.

Table S3: Comparing the AUROC for transductive learning setting in K562 cell line with varying percent of edges in the test set (validation set contains 10% edges in all cases). ± denotes the standard deviation on 10 independent trials.

| Test | RNAcommender | GCN | W | RNA | W.RNA | SN | W.SN | RNA.SN | W.RNA.SN | SN.RL | W.SN.RL | RNA.SN.RL | W.RNA.SN.RL |
|---|---|---|---|---|---|---|---|---|---|---|---|---|---|
| 10% | 0.604 ±0.003 | 0.751 ±0.002 | 0.723 ±0.003 | 0.768 ±0.004 | 0.739 ±0.003 | 0.740 ±0.004 | 0.731 ±0.004 | 0.759 ±0.003 | 0.742 ±0.002 | 0.751 ±0.002 | 0.740 ±0.004 | 0.758 ±0.004 | 0.742 ±0.004 |
| 20% | 0.601 ±0.004 | 0.736 ±0.002 | 0.715 ±0.004 | 0.757 ±0.002 | 0.731 ±0.003 | 0.729 ±0.003 | 0.719 ±0.004 | 0.755 ±0.002 | 0.734 ±0.004 | 0.738 ±0.002 | 0.726 ±0.003 | 0.750 ±0.003 | 0.730 ±0.004 |
| 30% | 0.595 ±0.004 | 0.717 ±0.002 | 0.700 ±0.002 | 0.740 ±0.002 | 0.719 ±0.003 | 0.714 ±0.004 | 0.705 ±0.003 | 0.739 ±0.002 | 0.722 ±0.003 | 0.720 ±0.002 | 0.708 ±0.002 | 0.732 ±0.003 | 0.714 ±0.003 |
| 40% | 0.594 ±0.005 | 0.717 ±0.003 | 0.699 ±0.004 | 0.740 ±0.001 | 0.718 ±0.003 | 0.715 ±0.001 | 0.703 ±0.004 | 0.739 ±0.002 | 0.720 ±0.003 | 0.720 ±0.002 | 0.707 ±0.002 | 0.731 ±0.002 | 0.711 ±0.002 |
| 50% | 0.586 ±0.005 | 0.709 ±0.003 | 0.690 ±0.002 | 0.730 ±0.002 | 0.708 ±0.002 | 0.707 ±0.002 | 0.694 ±0.002 | 0.730 ±0.001 | 0.712 ±0.002 | 0.711 ±0.001 | 0.695 ±0.002 | 0.722 ±0.002 | 0.702 ±0.003 |

Table S4: Comparing the AP for transductive learning setting in K562 cell line with varying percent of edges in the test set (validation set contains 10% edges in all cases). ± denotes the standard deviation on 10 independent trials.

| Test | RNAcommender | GCN | W | RNA | W.RNA | SN | W.SN | RNA.SN | W.RNA.SN | SN.RL | W.SN.RL | RNA.SN.RL | W.RNA.SN.RL |
|---|---|---|---|---|---|---|---|---|---|---|---|---|---|
| 10% | 0.607 | 0.754 | 0.728 | 0.774 | 0.747 | 0.751 | 0.739 | 0.775 | 0.754 | 0.744 | 0.734 | 0.741 | 0.729 |
| | ±0.004 | ±0.002 | ±0.004 | ±0.004 | ±0.004 | ±0.003 | ±0.004 | ±0.002 | ±0.002 | ±0.002 | ±0.005 | ±0.004 | ±0.007 |
| 20% | 0.604 | 0.740 | 0.720 | 0.765 | 0.738 | 0.738 | 0.729 | 0.769 | 0.747 | 0.732 | 0.722 | 0.733 | 0.718 |
| | ±0.006 | ±0.003 | ±0.003 | ±0.003 | ±0.003 | ±0.003 | ±0.004 | ±0.002 | ±0.004 | ±0.003 | ±0.003 | ±0.004 | ±0.006 |
| 30% | 0.600 | 0.725 | 0.707 | 0.751 | 0.727 | 0.724 | 0.715 | 0.756 | 0.735 | 0.716 | 0.706 | 0.719 | 0.703 |
| | ±0.005 | ±0.002 | ±0.002 | ±0.003 | ±0.003 | ±0.003 | ±0.003 | ±0.002 | ±0.002 | ±0.002 | ±0.004 | ±0.006 | ±0.004 |
| 40% | 0.598 | 0.721 | 0.705 | 0.749 | 0.725 | 0.720 | 0.709 | 0.750 | 0.730 | 0.713 | 0.701 | 0.714 | 0.698 |
| | ±0.008 | ±0.003 | ±0.004 | ±0.001 | ±0.003 | ±0.002 | ±0.003 | ±0.002 | ±0.003 | ±0.002 | ±0.003 | ±0.004 | ±0.003 |
| 50% | 0.586 | 0.710 | 0.693 | 0.735 | 0.714 | 0.709 | 0.697 | 0.738 | 0.719 | 0.702 | 0.688 | 0.703 | 0.687 |
| | ±0.008 | ±0.003 | ±0.003 | ±0.003 | ±0.002 | ±0.003 | ±0.003 | ±0.002 | ±0.002 | ±0.003 | ±0.003 | ±0.002 | ±0.004 |

In Section 4.1.2, we described the feature extraction method used for the experiments in the main text. Features can also be extracted by treating RNA and protein sequences as a special kind of language, where k-mers can be treated as words and sequences as sentences. Natural language processing techniques such as word2vec (Mikolov *et al.*, 2013) can then be used to learn embeddings for protein and RNA sequences (Asgari and Mofrad, 2015). Results for this alternative feature extraction scenario for can be seen in Table S5. While the vanilla GCN is only a little better than a random classifier, we observe that appending RNA-seq to the final embeddings significantly improves the performance (with and without structured negative sampling). This further highlights the importance of RNA-seq data.

Table S5: Comparing the AUROC for transductive learning setting in K562 cell line when *word2vec-based node features* are used while varying the percent of edges in the test set (validation set contains 10% edges in all cases). The bold marker denotes the best performing model(s) based on a t-test. The error bar ± denotes the standard deviation of the test performance of 10 independent trials.

| Test | GCN | RNA | RNA.SN |
|---|---|---|---|
| 10% | 0.576±0.003 | **0.769**±0.002 | 0.762±0.003 |
| 20% | 0.573±0.004 | **0.758**±0.002 | 0.752±0.002 |
| 30% | 0.563±0.004 | **0.741**±0.002 | **0.739**±0.002 |
| 40% | 0.584±0.002 | **0.740**±0.002 | 0.738±0.002 |
| 50% | 0.593±0.002 | **0.729**±0.002 | **0.729**±0.002 |

In Figure S2 we track the performance of the two best performing GNN models, GCN and GCN (RNA), as the percent of edges in the test set are increased (or equivalently the model is trained on fewer interactions) from 40% to 80%. We observe that even when the test set contains 80% of the interactions, the models achieve an AUC of greater than 0.6. The plot also shows that performance starts to degrade more rapidly when 60% to 80% of interactions are in the test set.

We also show the results for inductive link prediction in the HepG2 cell line in Figure S3, which compares the performance of different models over the entire set of proteins. Each box plots the distribution of mean AUROC/AP for proteins in the HepG2 cell line (10 replications for each protein). Again, we observe that

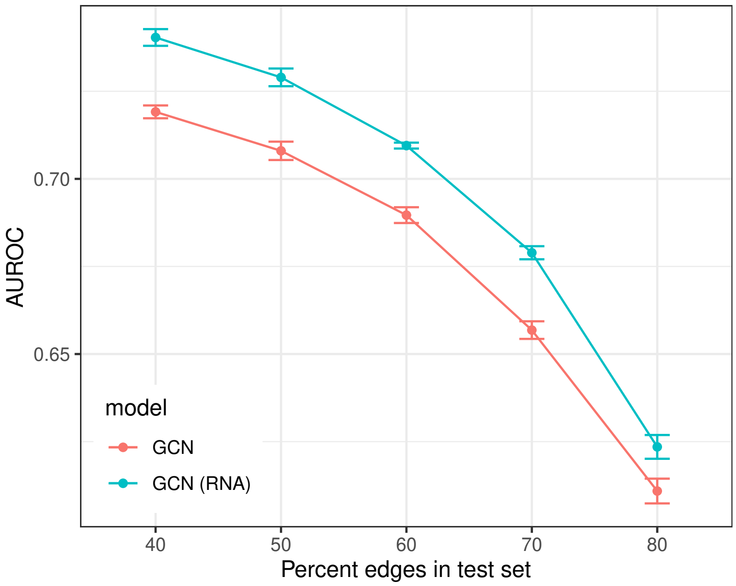


Figure S2: The performance of the two GCN settings as we increase the percentage of edges in the test set. The error bars show the standard deviation on 10 independent trials.

the choice of similarity function has very little impact, but unlike for K562, appending RNA-seq to the final embeddings does improve model performance.

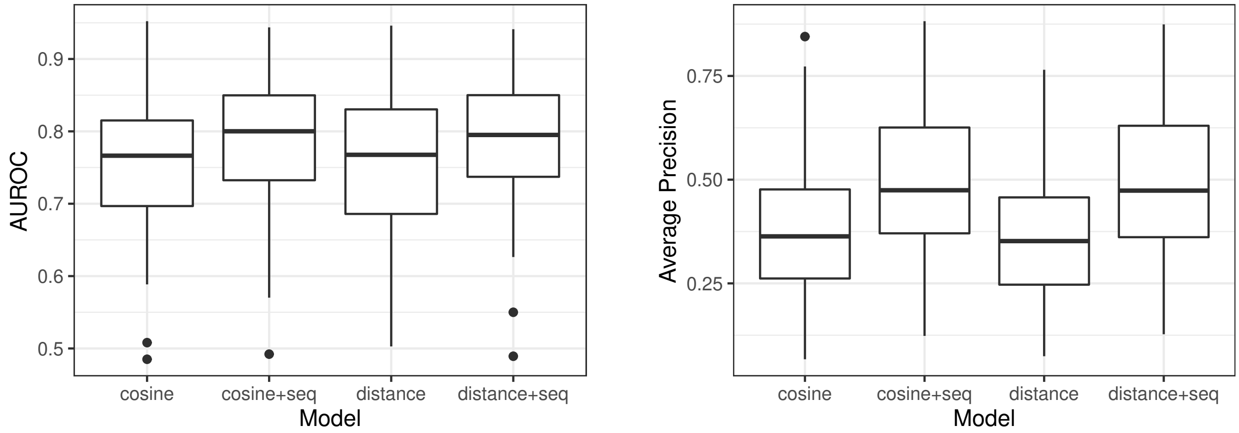


Figure S3: Comparing the performance of various models for *de novo* prediction in HepG2 cell line. Each box shows the distribution of mean AUROC (left) or average precision (right) over the entire set of proteins when the model is tested for a single protein in the test set.

Finally, we perform a comparison with NPI-GNN (Shen *et al.*, 2021) on the smaller datasets used in their paper. We would like to point out that while both papers use GNNs for predicting RNA-protein interactions, the architectures are very different. NPI-GNN has approximately 30 times more parameters than our approach (153154 vs 5610), leading to substantially larger memory and computational requirements.

For instance, NPI-GNN could not process our data even with 200GB of memory on a supercomputing server; by comparison, our model's total memory footprint on the same data is 3GB. Preliminary results (without hyperparameter tuning) in Table S6 show the performance of our model on the datasets used by NPI-GNN to be competitive, with a moderate loss in performance for the massive computational gain.

Table S6: Comparing performance of our approach with NPI-GNN on the datasets in (Shen *et al.*, 2021).

| Dataset | Method | AUC | Accuracy | Sensitivity | Specificity | Precision | MCC |
|---|---|---|---|---|---|---|---|
| NPInter2.0 | IPMiner | - | 0.952 | 0.905 | 0.959 | 0.895 | 0.896 |
| | NPI-GNN | 0.970 | 0.933 | 0.956 | 0.911 | 0.915 | 0.868 |
| | Ours | 0.951 | 0.897 | 0.894 | 0.901 | 0.892 | 0.794 |
| RPI7317 | IPMiner | - | 0.913 | 0.902 | 0.924 | 0.922 | 0.827 |
| | NPI-GNN | - | 0.915 | 0.927 | 0.907 | 0.907 | 0.830 |
| | Ours | 0.917 | 0.865 | 0.874 | 0.854 | 0.876 | 0.731 |
